# Microrna-184 is Induced by Store-Operated Calcium Entry and Regulates Early Keratinocyte Differentiation

**DOI:** 10.1101/319541

**Authors:** Adam Richardson, Andrew Powell, Darren W. Sexton, Jason L. Parsons, Nicholas J. Reynolds, Kehinde Ross

## Abstract

Extracellular calcium (Ca^2+^) and store-operated Ca^2+^ entry (SOCE) govern homeostasis in the mammalian epidermis. Multiple microRNAs (miRNA) also regulate epidermal differentiation, and raised external Ca^2+^ modulates the expression of several such miRNAs in keratinocytes. However, little is known about the regulation of miR-184 in keratinocytes or the roles of miR-184 in keratinocyte differentiation. Here we report exogenous Ca^2+^ stimulates miR-184 expression in primary epidermal keratinocytes and that this occurs in a SOCE-dependent manner. Levels of miR-184 were raised by about 30-fold after exposure to 1.5 mM Ca^2+^ for 5 days. In contrast, neither phorbol ester nor 1, 25-dihydroxyvitamin D3 had any effect on miR-184 levels. Pharmacologic and genetic inhibitors of SOCE abrogated Ca^2+^-dependent miR-184 induction by 70% or more. Ectopic miR-184 inhibited keratinocyte proliferation and led to a 4-fold increase in the expression of involucrin, a marker of early keratinocyte differentiation. Exogenous miR-184 also triggered a 3-fold rise in levels of cyclin E and doubled the levels of γH2AX, a marker of DNA double strand breaks. The p21 cyclin-dependent kinase (CDK) inhibitor, which supports keratinocyte growth arrest, was also induced by miR-184. Together our findings point to a SOCE:miR-184 pathway that targets a cyclin E/DNA damage regulatory node to facilitate keratinocyte differentiation.

## INTRODUCTION

The epidermis is a stratified tissue maintained by controlled proliferation and terminal differentiation of keratinocytes (Eckhart et al., 2013). Extracellular calcium (Ca^2+^) induces involucrin (IVL) and several other differentiation proteins to enable formation of the cornified envelope. Differentiating suprabasal keratinocytes express cell cycle proteins such as cyclin A, B and E to support their enlargement (Zanet et al., 2010). Furthermore, cyclin E, which drives the G1/S phase transition, accumulates within IVL-expressing suprabasal layers and promotes the proliferation:differentiation switch through mitotic failure and DNA damage (Freije et al., 2012, Zanet et al., 2010).

MicroRNAs (miRNAs) are short noncoding RNAs (18–25 nucleotides) that attenuate post-transcriptional gene output through translational inhibition and destabilization of mRNA transcripts, the latter appearing to sustain the bulk of steady-state repression (Eichhorn et al., 2014, Huntzinger and Izaurralde, 2011). Several miRNAs have been implicated in epidermal differentiation including miR-203, miR-205 and miR-24 (reviewed (Riemondy et al., 2014)). Furthermore, over 50 miRNAs were upregulated in human keratinocytes exposed to high extracellular Ca^2+^ (Hildebrand et al., 2011) and miR-24 was shown to be upregulated by Ca^2+^-during keratinocyte differentiation (Amelio et al., 2012). Given that store-operated Ca^2+^ entry (SOCE) through the STIM1:ORAI1 axis plays essential roles in keratinocyte and epidermal physiology (Numaga-Tomita and Putney, 2013, Ross et al., 2007, Vandenberghe et al., 2013), it is conceivable that Ca^2+^-dependent induction of miRNAs relies at least partly on SOCE. However, the impact of SOCE on miRNA expression has received little attention.

Very recent studies detected miR-184 expression predominantly in the spinous layer of neonatal mouse epidermis, thus implicating miR-184 in epidermal differentiation (Nagosa et al., 2017). Modulation of miR-184 levels in mouse skin and cultured human keratinocytes together revealed that miR-184 represses keratinocyte proliferation and supports commitment to differentiation through the Notch axis. Although not listed among miRNAs upregulated by high Ca^2+^ in earlier work (Hildebrand et al., 2011), a 4-fold increase in miR-184 expression was observed in human keratinocytes exposed to high Ca^2+^ for 7 days (Nagosa et al., 2017). However, the mechanisms of miR-184 induction have not been defined and little is known about the signalling pathways regulating miR-184 expression.

We previously observed expression of miR-184 in reconstituted human epidermis (RHE) and in the HaCaT keratinocyte cell line (Roberts et al., 2013). In contrast, Lavker and colleagues did not detect miR-184 in monolayer cultures of proliferating epidermal keratinocytes (Yu et al., 2008). Exposure of RHE to inflammatory cytokines interleukin-22 (IL-22) or oncostatin M (OSM) enhanced miR-184 expression in our studies, suggesting that miR-184 levels can be modulated by external signals (Roberts et al., 2013). Here, we show that elevation of extracellular Ca^2+^ induces miR-184 in human primary epidermal keratinocytes (HPEK) in a monolayer culture. The upregulation of miR-184 was associated specifically with Ca^2+^ as other differentiation stimuli did not trigger miR-184 expression. Furthermore, pharmacologic or genetic inhibition of SOCE impaired the induction of miR-184. Finally, we found that ectopic miR-184 itself promoted HPEK differentiation and this was associated with the elevation of cyclin E, DNA damage and induction of p21.

## RESULTS AND DISCUSSION

### Induction of miR-184 in Human Primary Epidermal Keratinocytes

We evaluated miR-184 expression in HPEK maintained in parallel cultures under low Ca^2+^ (0.07 mM) or high (1.5 mM) extracellular Ca^2+^ to sustain proliferation or promote differentiation, respectively. As shown in Fig 1a, relative miR-184 expression was almost 30-fold higher in HPEK treated with 1.5 mM Ca^2+^ for 5 days compared to cells maintained in low Ca^2+^. Only very low levels of miR-184 were detected in the cells under low Ca^2+^ conditions, suggesting that the high Ca^2+^ challenge triggers *de novo* miR-184 expression. In contrast, the active form of vitamin D, 1, 25-dihydroxyvitamin D_3_, (1, 25-(OH)_2_D3; Calcitriol) or phorbol 12-myristate 13-acetate (PMA) did not promote miR-184 expression in HPEK over the time points assessed (Fig.1b,c). By comparison, all three agents evoked robust *IVL* expression as expected (Fig.1d-f) given their established functions as inducers of keratinocyte differentiation (Bikle, 2004, Karlsson et al., 2010). Together, these observations suggest that the induction of miR-184 by Ca^2+^ is associated with Ca^2+^-dependent pathways and not simply keratinocyte differentiation. Interferon gamma (IFNγ) has also been reported to trigger keratinocyte differentiation (Karlsson et al., 2010) but no induction of miR-184 was observed with 10 ng/ml IFNγ, and the low levels of miR-184 registered in untreated cells became undetectable following IFNγ stimulation for 1 or 5 days (data not shown).

**Figure 1:**
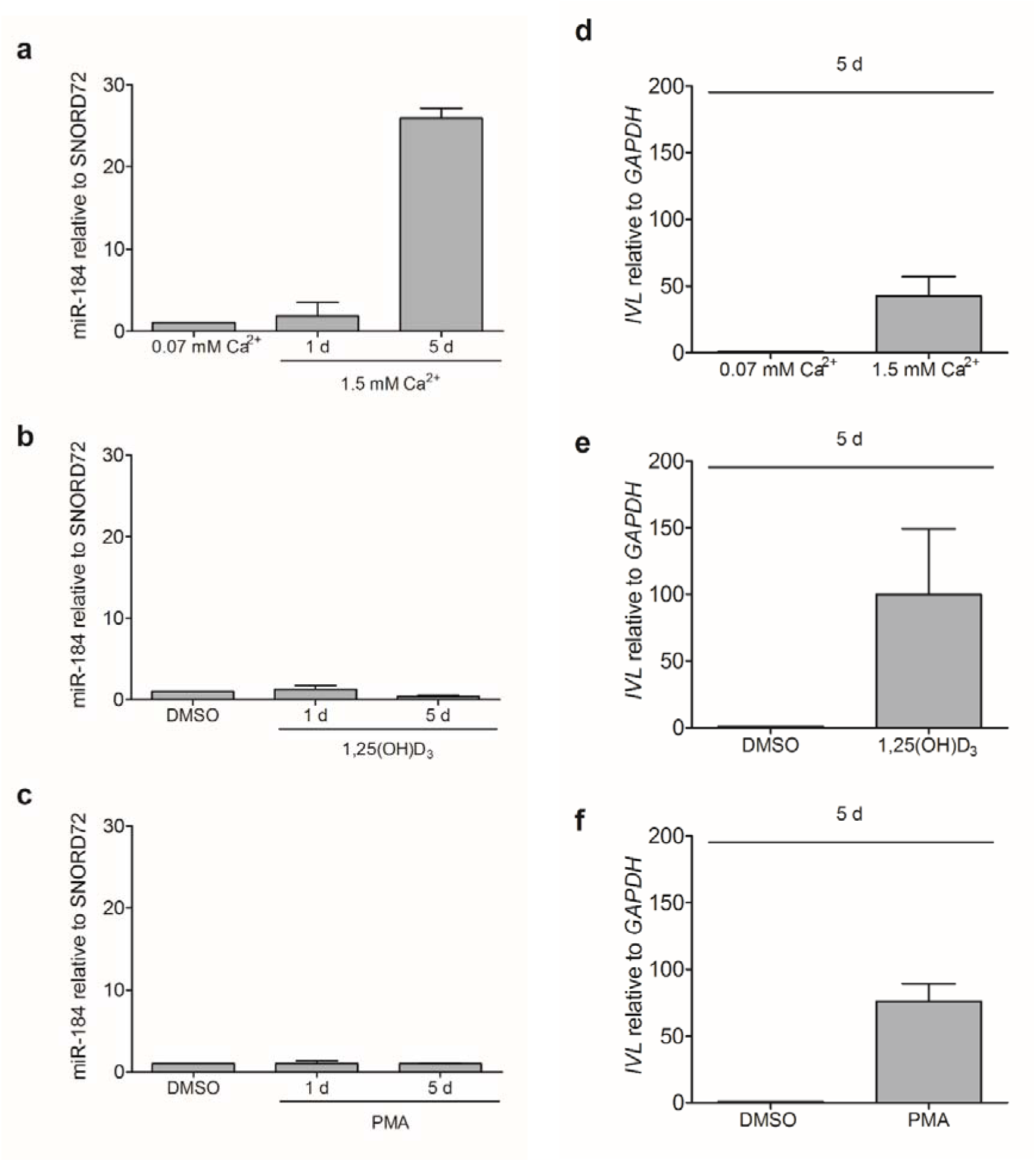
Induction of miR-184 during Ca^2+^-dependent HPEK differentiation. (a) HPEKs grown to confluence were treated with high (1.5 mM) Ca^2+^, 100 nM 1,25(OH)D_3_, 100 nM PMA or dimethyl sulfoxide (DMSO) vehicle for 1 or 5 days (d). (b) Upregulation of the HPEK differentiation marker involucrin *(IVL)* during 5 d exposure to 1.5 mM Ca^2+^, 100 nM 1,25(OH)D_3_ or 100 nM PMA. Data shown represent means +SEM from 3 independent experiments. Expression was normalized to SNORD72 for miR-184 and *GAPDH* for *IVL*. Values are presented relative to 0.07 mM Ca^2+^ or DMSO-treated controls.

To determine whether Ca^2+^-dependent induction of miR-184 was associated with SOCE, HPEK were maintained in high Ca^2+^ for 5 days in the presence of the SOCE blocker gadolinium (III) Gd^3^+ or the pharmacologic ORAI1 inhibitor BTP2. Alternatively, ORAI1 expression was ablated using short-interfering RNA (siRNA) that was transferred into the cells using nucleofection 1 d prior to Ca^2+^ elevation. We observed that Gd^3^+, BTP2 and siORAI1 reduced Ca^2+^-induced miR-184 expression by 70% or more (Fig.2a-c). The siRNA suppressed ORAI1 levels by about 55%, whereas Gd^3^+ and BTP2 had little effect on ORAI1 expression (Supplementary Fig. S1). Together, these findings suggest elevation of miR-184 during Ca^2+^-dependent keratinocyte differentiation is mediated at least partly by Ca^2+^ influx through ORAI1.

**Figure 2:**
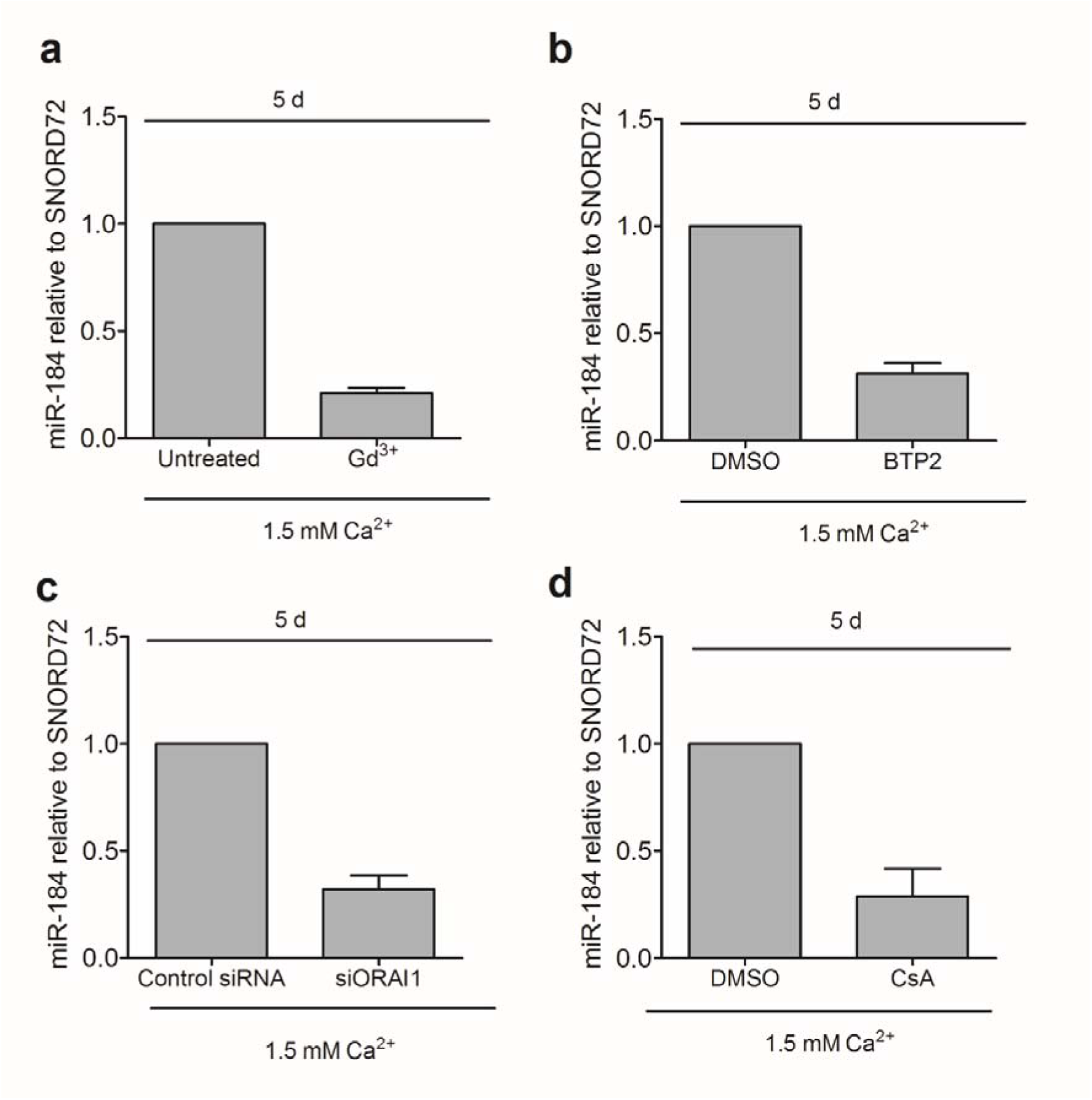
Inhibition of SOCE attenuates Ca^2+^-dependent miR-184 expression. HPEKs were maintained in 1.5 mM Ca^2+^ for 5 d with or without (a) 1 μM Gd^3^+, (b) 1 μM BTP2, (c) 100 nM ORAI1-targeting siRNA or (d) 1 μM CsA as indicated. The Gd^3^+, BTP2 and CsA were added 1 h prior to Ca^2+^ switch and refreshed after day 2. Data shown represent means +SEM from 3 independent experiments. Expression was normalized to SNORD72 for miR-184 and *GAPDH* for *IVL*. Values are presented relative to untreated, DMSO-treated or control siRNA as indicated.

The Ca^2+-^calmodulin/calcineurin-nuclear factor of activated T cells (NFAT) pathway is a prototypical effector of SOCE (Srikanth and Gwack, 2013). To examine the putative role of NFAT in miR-184 induction, HPEK were incubated with the indirect calcineurin inhibitor cyclosporin A (CsA) prior to high Ca^2+^ challenge. Treatment with 1 μΜ CsA inhibited Ca^2+^-dependent induction of miR-184 by 70% (Fig. 2d). Together, these observations indicates that SOCE mediates the induction of miR-184 in HPEK and this occurs at least partly through the Ca^2+-^calmodulin/calcineurin-NFAT axis.

### Effects of miR-184 on Keratinocyte Growth

The functional impact of miR-184 on HPEK behaviour was assessed using the MTT (3-(4,5-dimethylthiazol-2-yl)-2,5-diphenyltetrazolium bromide) reduction assay. Metabolic activity was reduced by around 40% in proliferating HPEK transfected with a miR-184 mimic compared control cells (Fig. 3a) although this was not significant. By comparison, live/dead cell staining with the Trypan Blue exclusion assay indicated a 17% drop in cell viability (Fig. 3b). Cell cycle analyses revealed that the proportion of HPEKs in G1 rose from 63.6% in controls to 72.7% in cells transfected with the miR-184 (Fig. 3c and Supplementary Fig. S2). A corresponding drop in the percentage of cells in G2 was also observed, from 13.7% in controls to 5.7% in miR-184. The proportion of HPEKs in S phase remained relatively unchanged. Hence, miR-184 appears to reduce keratinocyte proliferation by inhibiting progression through the G1 phase of the cell cycle. These observations are consistent with the findings recently reported by Shalom-Feuerstein who showed that overexpression of miR-184 reduced keratinocyte proliferation in the basal layers of mouse epidermis (Nagosa et al., 2017).

**Figure 3:**
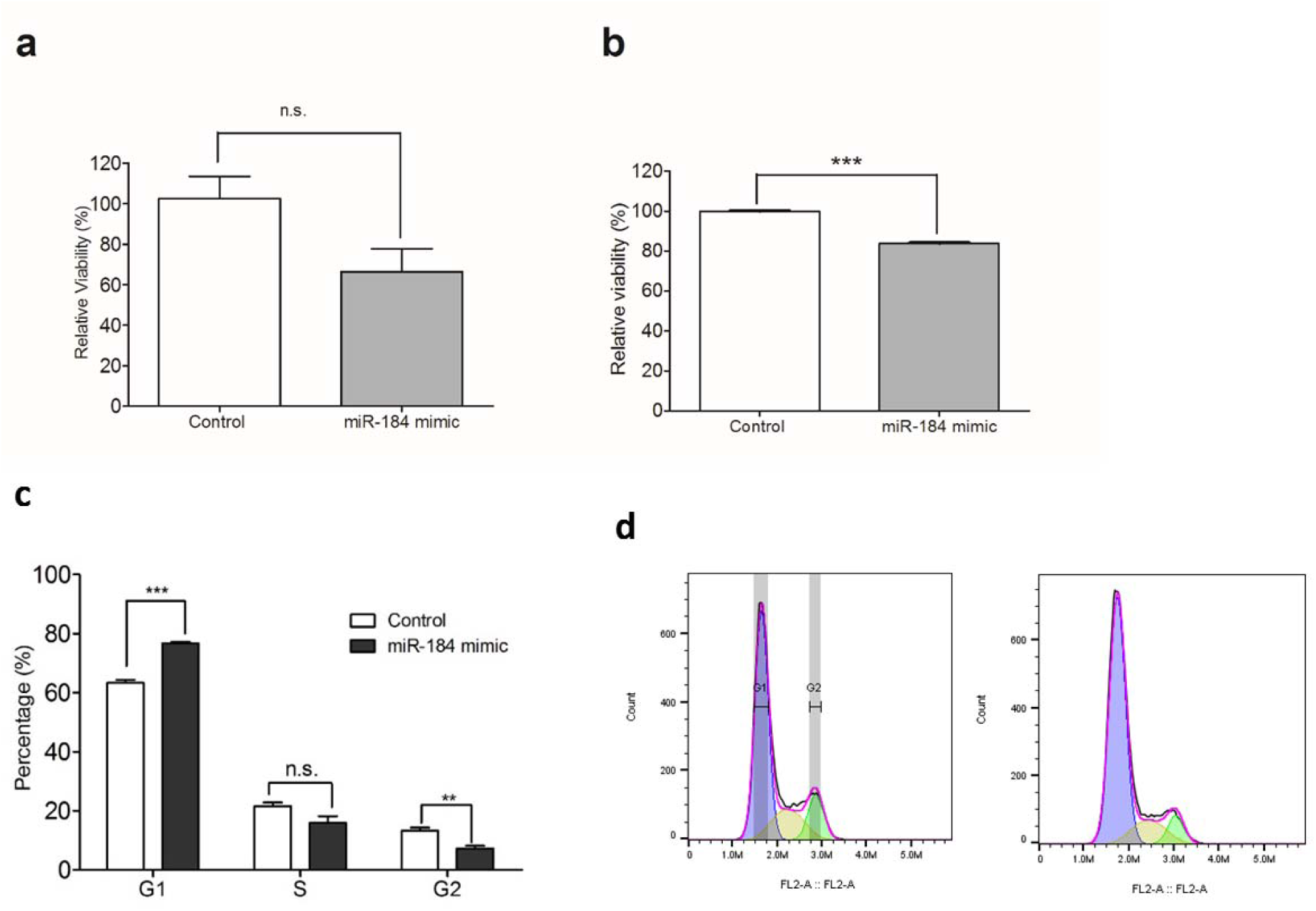
miR-184 reduces HPEK proliferation. HPEK were nucleofected with 100 nM of the miR-184 mimic or control oligonucleotide as indicated. The effect of miR-184 on HPEK viability determined with MTT (a) or trypan blue staining (b). Data shown are means +SEM of 3 independent experiments. (c) Cell cycle profiles were assessed using propidium iodide staining and flow cytometry. Representative histograms from HPEK loaded with miR-184 mimic *(left)* or control oligonucleotide *(right)* are presented in (d). ***, p<0.001; **, p<0.01; n.s., not significant.

### Effects of miR-184 on Keratinocyte Differentiation

To uncover the effects of miR-184 on keratinocyte differentiation, IVL and cyclin E levels were examined in HPEK transfected with the miR-184 mimic. Both proteins were barely detectable in proliferating HPEK maintained in low Ca^2+^ (Fig. 4a). In contrast, ectopic miR-184 triggered a 4-fold and 3-fold elevation in IVL and cyclin E expression, respectively (Fig. 4a,b). Conversely, both IVL and cyclin E were readily detected in HPEK cultured under high Ca^2+^ conditions. Furthermore. transfection of a locked nucleic acid (LNA) miR-184 inhibitor lowered IVL and cyclin E expression by 60% and 90% respectively (Fig. 4c,d). Taken together, these observations implicate miR-184 in the regulation of the cyclin E pathway proposed by Gandarillas and colleagues in which cyclin E hyperactivation causes DNA damage that in turn signals for growth arrest and subsequent keratinocyte differentiation (Freije et al., 2012, Zanet et al., 2010).

**Figure 4:**
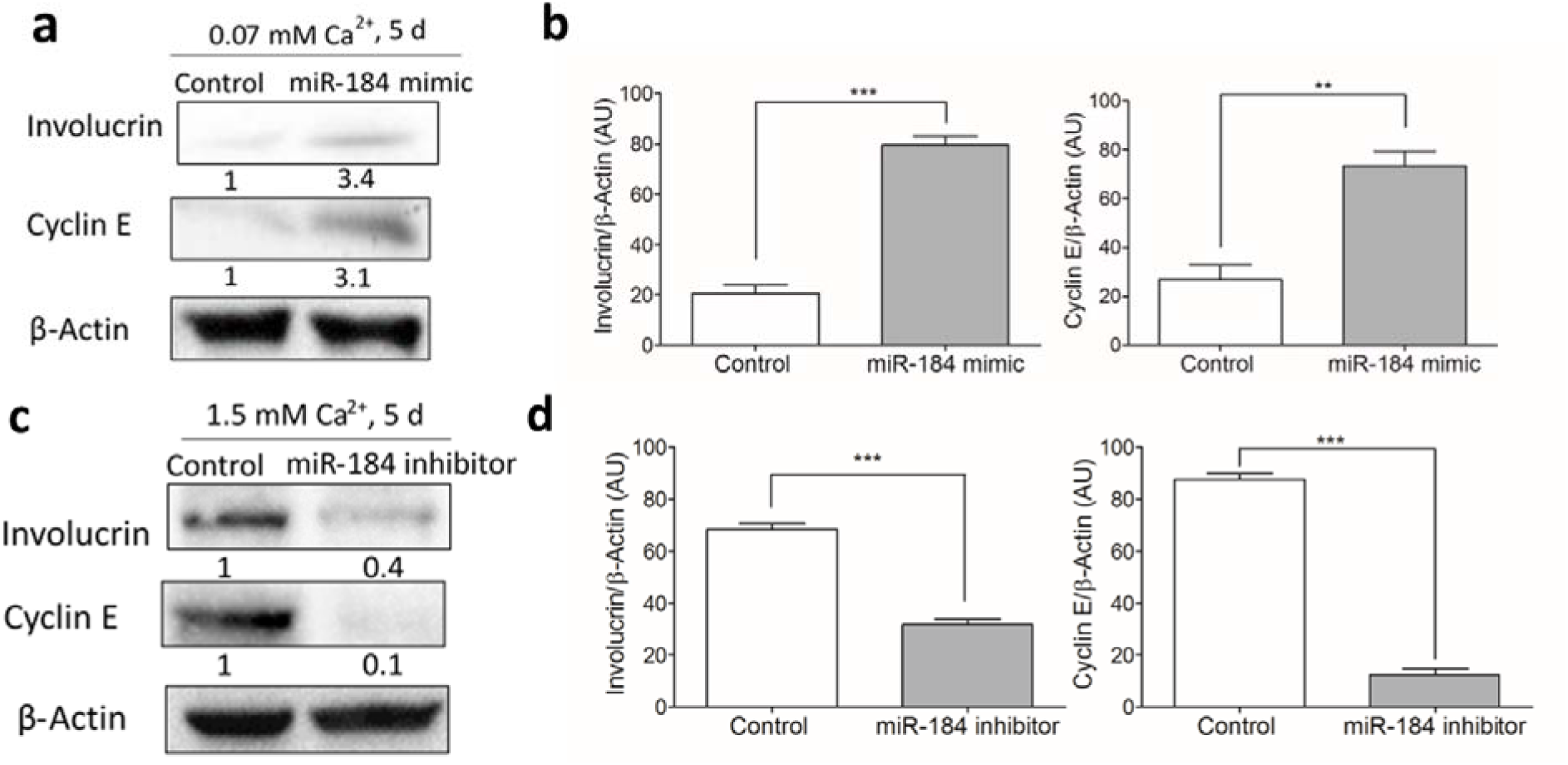
miR-184 induces IVL and cyclin E expression in HPEK. Cells loaded with 100 nM miR-184 mimic or a control oligonucleotide were maintained in low Ca^2+^ (0.07 mM) media (a, b) or high Ca^2+^ (1.5 mM) media (c,d) for 5 d prior to western blotting. Graphs (b, d) show the mean + SEM densitometry levels relative to β-actin. Data shown were pooled from 3 independent experiments. ***, p<0.001; **, p<0.01.

### Induction of DNA damage and p21 by miR-184

Given the observed miR-184–dependent induction of cyclin E (Fig. 4a,b) and the ability of cyclin E to promote DNA damage in HPEK (Freije et al., 2012, Zanet et al., 2010), we hypothesised that ectopic miR-184 would trigger induction of DNA damage. Levels of γH2AX, a biomarker of DNA double-strand breaks, were elevated almost 2-fold in HPEK transfected with miR-184 mimic compared to their counterparts transfected with a control mimic (Fig. 5a). Furthermore, transfection with miR-184 mimic doubled the number of γH2AX foci when visualised by fluorescence microscopy (Fig. 5b). Finally, we investigated the effect of miR-184 on the p21-CDK inhibitor, which previous studies showed was upregulated by cyclin E in HPEK (Freije et al., 2012). As shown in Fig. 5c,d exogenous miR-184 mimic increased p21 expression by 2.5-fold at the transcript level and 2-fold at the protein level.

Taken together, our findings uncover a mechanism whereby SOCE triggers miR-184 during keratinocyte differentiation. In turn, miR-184 evokes DNA damage and p21 expression, the latter of which may explain the ability of miR-184 to inhibit keratinocyte progression through G1 phase of the cell cycle (Fig.3c). Questions remain though, about the mechanisms by which accentuation of miR-184 promotes DNA damage. The model of Gandarillas and colleagues (Freije et al., 2012, Zanet et al., 2010) would suggest that elevation of cyclin E drives DNA damage but we have not addressed the impact of cyclin E on miR-184-dependent DNA damage in the present work. Assuming such a model is valid, there is still the question of how cyclin E is upregulated by miR-184 since no miR-184 binding sites were predicted in *CCNE1 or CCNE2* transcripts using TargetScan 7.1. Hence the genomic or transcriptomic targets that convert miR-184 binding into a DNA damage signal remain to be defined. Nonetheless, we propose that in addition to targeting the Notch-dependent pathway (Nagosa et al., 2017), miR-184 serves to couple SOCE to the induction of cyclin E and DNA damage in order to elevate p21 and involucrin expression (Fig. 5e). Determining whether other miRNAs associated with keratinocyte differentiation exploit the cyclin E:DNA damage pathway will reveal whether this is a common mechanism or one unique to miR-184 for to sustaining epidermal homeostasis.

**Figure 5:**
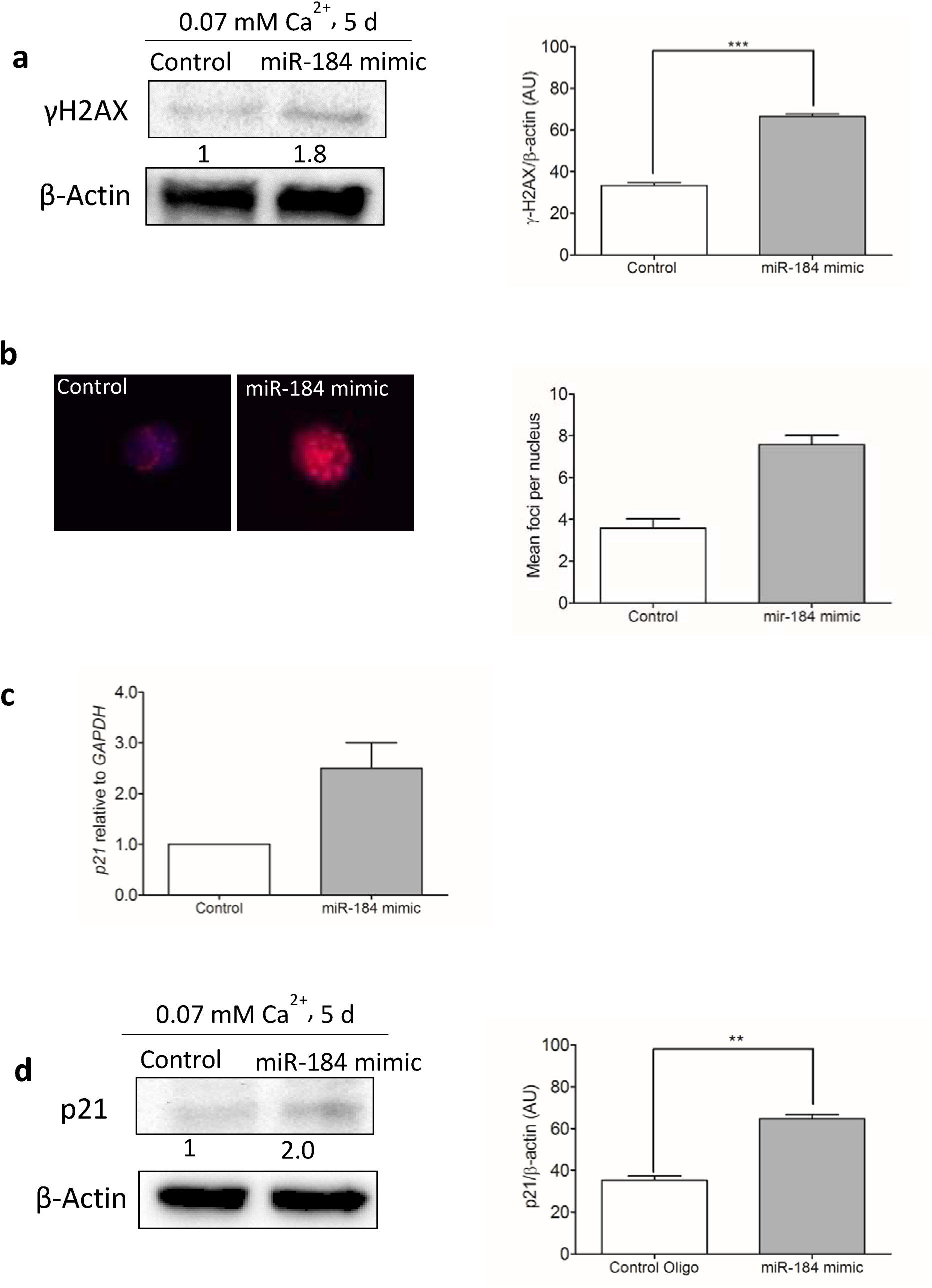
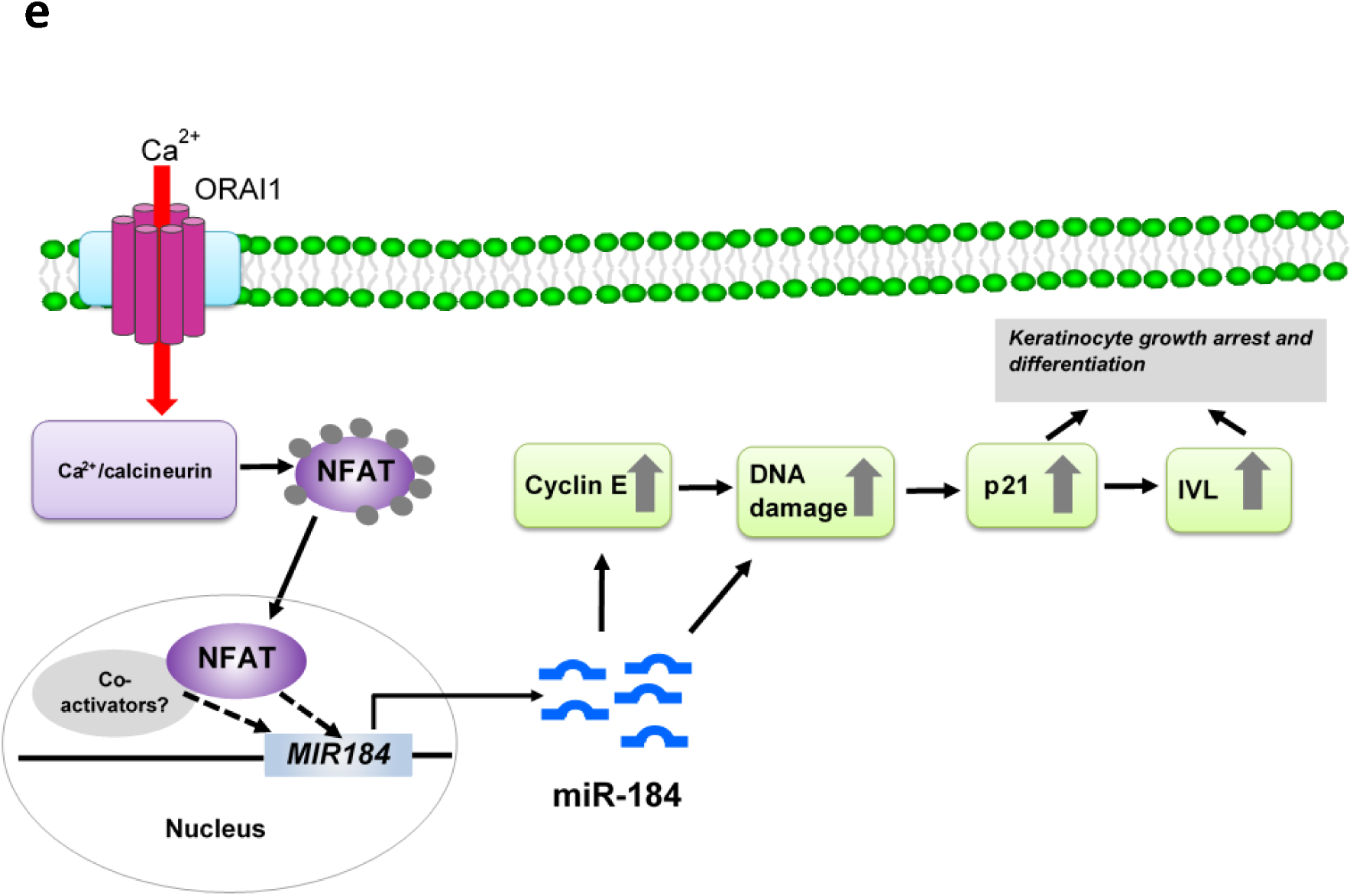
miR-184 promotes DNA damage and p21 expression in HPEK. (a) Cells loaded with 100 nM miR-184 mimic or a control oligonucleotide were maintained in low Ca^2+^ (0.07 mM) media for 5 d prior to western blotting for γH2AX. (b) Overlaid immunofluorescent images for γH2AX and DAPI staining in cells loaded with 100 nM miR-184 for 5 d. The graph depicts the foci number. (c, d) Elevated expression of p21 at the transcript (c) and protein (d) levels in HPEK loaded with 100 nM miR-184 for 5 d. Graphs in (a), (b) and (d) show the mean + SEM densitometry levels relative to β-actin, pooled from three independent experiments. ***, p<0.0001; **, p<0.001. (e) Schematic representation of the SOCE:miR-184 axis to keratinocyte differentiation.

## METHODS

### Reagents

Oligonucleotides (miR-184 mimic, siORAI and respective controls) were purchased from GE Healthcare (Little Chalfont, UK). The locked nucleic acid (LNA) miR-184 inhibitor and a non-targeting control were from Exiqon (Vedbaek, Denmark). Gadolinium(III) chloride (Gd^3+^) and differentiation reagents (1, 25-(OH)_2_D3/calcitriol and PMA) and were purchased from Bio-Techne (Abingdon, UK). BTP2 (also known as YM58483 was purchased from Abcam (Cambridge, UK).

### Keratinocyte Isolation, Culture and Differentiation

Human progenitor epidermal keratinocytes (HPEK) were isolated from human foreskin or purchased from CellnTec (Bern, Switzerland). For isolation, tissues were washed with phosphate-buffered saline, trimmed of any subcutaneous fat, connective tissues and blood vessels before digestion with 20 mg/ml dispase (Sigma) at 4°C overnight with further digestion for 1 hour at room temperature. Thereafter, the epidermis was separated and incubated in 0.5% TrpLE (Thermofisher scientific, Cheshire, UK) for 5 min at 37°C, 5% CO_2_. Keratinocytes were centrifuged (1200 rpm for 5 min) and re-suspended in CnT-Prime (CellnTec) supplemented with 1% penicillin/streptomycin/amphotericin B (PSA) and IsoBoost supplement CnT-ISO (CellnTec) to enhance isolation efficiency. Culture medium was changed every 2–3 days until cells reached 80% confluence with PSA exclusion from the culture medium after the first passage. Keratinocytes were sub-cultured using CnT-Accutase (CellnTec) and re-cultured at 4 × 10^3^ cells per cm^2^. For differentiation, keratinocytes were at 3 × 10^5^/well of a 6-well plate in CnT-Prime and allowed to reach confluence before adjusting the medium to 1.5 mM calcium using CaCl_2_.

### Oligonucleotide Nucleofection

Keratinocytes were sub-cultured and 5 × 10^5^ cells resuspended in 100 μl nucleofection solution from the P3 Primary Cell 4D-Nucleofector kit (Lonza, Castleford, UK) and with 100 nM human miR-184 mimic (a synthetic double-stranded oligonucleotide mimicking endogenous miR-184), 100 nM of LNA miR-184 inhibitor or respective non-targeting negative control oligonucleotides. Modulation of miR-184 levels by the mimic or inhibitor was confirmed by sqRT-PCR (Supplementary Fig. S3). Cell suspensions were then transferred to nucleofection cuvettes and pulsed on the DS-138 programme of a 4D-Nucleofector. After incubation in pre-equilibrated CnT-Prime at room temperature for 10 min, the nucleofected cells were transferred to multiwall plates and incubated at 37°C, 5% CO_2_ incubator with the media replacement the following day.

### Cell Viability

Cells transfected with the miR-184 mimic or negative control were seeded at 2×10^4^/well of a 96-well plate and maintained in CnT-Prime for 3 d. The MTT reagent 3-(4,5-Dimethylthiazol-2-yl)-2,5-diphenyltetrazolium bromide was added to each well at 5 mg/ml and incubated at 37°C, 5% CO_2_ for 4 h. Culture medium was removed, the 96-well plate air dried and 100 μl dimethyl sulfoxide (DMSO) added to each well. After shaking for 5 min to ensure solubilisation of formazan crystals, the absorbance of each well was read on a Clariostar plate reader (BMG Labtech, Aylesbury, UK) at OD 470 nm. All experiments were performed in triplicate and at least three times. For trypan blue viability tests, an aliquot of cells nucleofected with miR-184 mimic or negative control oligo was mixed 1:1 with 0.4% trypan blue before loading onto a haemocytometer. Dark blue cells were counted as non-viable cells and those with bright centres counted as live.

### Cell Cycle

Nucleofected cells seeded at 3 × 10^5^/well of a 6-well plate were grown for 2 d, detached using CnT-Accutase (CellnTec, Bern, Switzerland) washed twice with PBS then fixed in 70% ice cold ethanol for 24–72 h. Cells were washed with PBS once more and propidium iodide (100 μg/ml) was added for 30 min in the dark at room temperature. Cells were then examined using a BD Accuri C6 flow cytometer (BD Biosciences, Wokingham, UK) with gates for forward scatter vs side scatter (FSC vs SSC) to exclude debris and FSC height vs FSC area (FSC-H vs FSC-A) to discriminate against doublets. Representative gating plots are presented in Supplementary Fig. S2. Fluorescent values from 1 × 10^4^ gated events was collected using the FL-2A parameter. Analysis was performed using FlowJo version 10.0 software and univariant Dean-Jett-Fox algorithm.

### Semi-quantitative Reverse Transcriptase PCR (sqRT-PCR)

Total RNA was isolated from cells using the AllPrep DNA/RNA/miRNA Universal kit (Qiagen, Manchester, UK). RNA concentration was determined using a NanoDrop™ 2000c. Complementary DNA (cDNA) was synthesised from 400 ng of RNA using the miScript II RT kit (Qiagen) with HiFlex buffer. PCR amplifications were performed with Quantifast SYBR Green and QuantiTect miRNA/universal primers or RT^2^ mRNA primer assays, all from Qiagen. Thermocycling was performed on a Rotor-Gene^®^ as follows: 95°C for 15 min, followed by 40 cycles of 94°C for 15 s, 55°C for 30 s and 70°C for 30 s. Relative expression of miRNA and mRNA determined using the 2^−∆∆CT^ relative quantification method (Livak and Schmittgen, 2001). Cycle threshold values are presented in Supplementary Tables 1 and 2.

### Western Blotting

Cells were lysed in RIPA buffer containing protease and phosphatase inhibitors and 20 or 40 μg of total protein resolved on mini-PROTEAN 12% SDS/PAGE precast gels (Bio-Rad, Watford, UK). After transfer to polyvinylidene difluoride (PVDF) membranes, samples were incubated with primary antibodies: Involucrin (1:1000; Novus biologicals, Oxon, UK), Cyclin E (1:750; bio-Techne, Abingdon, UK), γH2AX (1:500; Millipore, Watford, UK), p21 (1:500; Bio-Rad, Oxfordshire, UK), GAPDH (1:5000, R&D systems) and β-actin (1:2000; Sigma) overnight at 4°C. Membranes were then washed several times with Tris-buffered saline with 0.1% Tween-20 (TBST) and incubated with horseradish peroxidase (HRP) conjugated secondary antibodies for 1 hour at room temperature. Membranes were washed before chemiluminescence was visualised using Clarity ECL reagents (Bio-Rad). ImageJ software was used to perform densitometry with target protein values normalised to the corresponding β-actin controls.

### Immunofluorescence staining for DNA double strand break foci

Immunostaining was performed as previously reported (Nickson et al., 2017). Briefly, fixed and peameabilised cells were incubated with 2% BSA in PBS with 0.1 % Tween-20 (PBST) for 1 h at room temperature to block non-specific staining. Incubation with γH2AX (1:1000) antibody was performed overnight at 4°C in PBST with 2% BSA. After washing, coverslips were incubated with goat anti-mouse Alexa Fluor 647 for 1 h at room temperature in the dark. Samples were then washed with PBS and mounted on a microscope slide using Fluoroshield containing DAPI (Sigma-Aldrich, Gillingham, UK). Cells were examined using a Leica DMI6000B fluorescent microscope supported by Leica Application Suite (LAS) X software.

### Statistics

Where indicated, statistical analysis was performed using the Students’ *t* test in Graph Pad Prism 5.0 (La Jolla, CA, USA).

## ACKNOWLEDGEMENTS

We thank Mr Abid Qazi, Mr Nadeem Haider, Dr Naseem Khan and Dr Mohammad Shafiq (Northern Circumcision Clinic, UK) for collecting skin samples (Liverpool John Moores University Research Ethics Committee approval number 16/PBS/008). We are grateful to Dr Eithne Costello (University of Liverpool, Liverpool, UK) for the gift of the cyclin E antibody. We thank Dr Katie Nickson and Dr Carlos Rubbi (both of the University of Liverpool, Liverpool, UK) for analysis and quantification of γH2AX foci staining. This study was funded by the British Skin Foundation (grant number 7006s). NJR's laboratory/research is supported by the NIHR-Newcastle Biomedical Research Centre and the Newcastle MRC/EPSRC Molecular Pathology Node.

